# Structural basis of light-induced redox regulation in the Calvin cycle

**DOI:** 10.1101/414334

**Authors:** Ciaran McFarlane, Nita R. Shah, Burak V. Kabasakal, Charles A.R. Cotton, Doryen Bubeck, James W. Murray

## Abstract

In plants, carbon dioxide is fixed via the Calvin cycle in a tightly regulated process. Key to this regulation is the conditionally disordered protein CP12. CP12 forms a complex with two Calvin cycle enzymes, glyceraldehyde-3-phosphate dehydrogenase (GAPDH) and phosphoribulokinase (PRK), inhibiting their activities. The mode of CP12 action was unknown. By solving crystal structures of CP12 bound to GAPDH, and the ternary GAPDH-CP12-PRK complex by electron cryo-microscopy, we reveal that formation of the N-terminal disulfide pre-orders CP12 prior to binding the PRK active site. We find that CP12 binding to GAPDH influences substrate accessibility of all GAPDH active sites in the binary and ternary inhibited complexes. Our model explains how CP12 integrates responses from both redox state and nicotinamide dinucleotide availability to regulate carbon fixation.

**One Sentence Summary:** How plants turn off carbon fixation in the dark.

---

The Calvin cycle fixes organic carbon to provide fuel for plants, a process tightly controlled by light-induced redox changes in regulated enzymes (*1*). Phosphoribulokinase (PRK) and glyceraldehyde-3-phosphate dehydrogenase (GAPDH) are essential enzymes in the photosynthetic dark reactions that control availability of substrate for the carboxylation enzyme Rubisco. PRK consumes ATP to produce the Rubisco substrate ribulose bisphosphate (RuBP), while GAPDH catalyses the reduction step of the Calvin cycle with NADPH to produce the sugar, glyceraldehyde 3-phosphate (GAP), which is used for regeneration of RuBP and is the main exit point of the cycle (*2*). GAPDH and PRK are co-regulated in response to the light reactions, which produce ATP and NADPH. In the light, NADPH and reduced ferredoxin are produced in the chloroplast stroma and the two disulfides of PRK are reduced to switch on PRK activity (*3*). GAPDH has no disulfides, instead redox regulation of GAPDH is driven by two disulfide bonds in the small regulatory inhibitor protein CP12 (*4, 5*), which is disordered under reducing conditions (*6*). In the dark, the stroma becomes oxidizing and intramolecular disulfide bonds within PRK and CP12 are formed. CP12 binds and inhibits GAPDH and then PRK activity in an obligate sequential reaction (*7*). CP12 is present in oxygenic phototrophs from cyanobacteria to plants (*8*). Previous structural studies of the GAPDH-CP12 complex resolved only a C-terminal fragment of CP12, so it remained unclear how CP12 could regulate both enzymes. A recent structure of a cyanobacterial CP12-CBS protein resolved an N-terminal CP12-like region, however the protein does not form a ternary complex with GAPDH and PRK (*9*). The lack of structural information for green-type PRK and CP12-bound regulatory complexes has prevented understanding a key mechanism of Calvin cycle redox control. To understand, at a molecular level, how the Calvin cycle is redox regulated in response to light, we solved crystal structures of a thermophilic cyanobacterial GAPDH with full-length CP12 and built an atomic model of the entire cyanobacterial GAPDH-CP12-PRK ternary complex using electron cryo-microscopy (cryoEM).

We co-crystallized recombinant *Thermosynechococcus elongatus* GAPDH and CP12 to obtain three crystal structures with density for full-length CP12, at a best resolution of 2.1 Å (table S1). The cyanobacterial GAPDH-CP12 complex has two molecules of CP12 per GAPDH tetramer (Fig. 1A). We call the two occupied sites proximal and the two unoccupied sites distal. In all our structures, and those published, the two proximal sites are in equivalent positions. As fully-occupied tetramers could be formed with excess CP12 (fig. S1), stoichiometry of the GAPDH-CP12-PRK complex remained unclear (*10, 11*). CP12 comprises an N-terminal PRK binding domain (residues 1-52) and a C-terminal GAPDH binding domain (residues 55-75) connected by a flexible linker (Fig. 1B). The N-terminal PRK binding domain is a two-helix bundle, stabilized by a disulfide bridge near the helical turn. The two helices bury a small hydrophobic core, including a short leucine zipper. Helix-2 (27-52) contains the conserved CP12 characteristic motif, AWDA(V/L)EEL (Fig. 1B, 2E), (*12*) which forms an acidic patch on the surface. Although the relative orientations of the N- and C-terminal domains of CP12 vary in our structures, the structures of both domains are conserved across four crystallographically independent molecules (Fig. 1C, 1D).

**Fig. 1.**
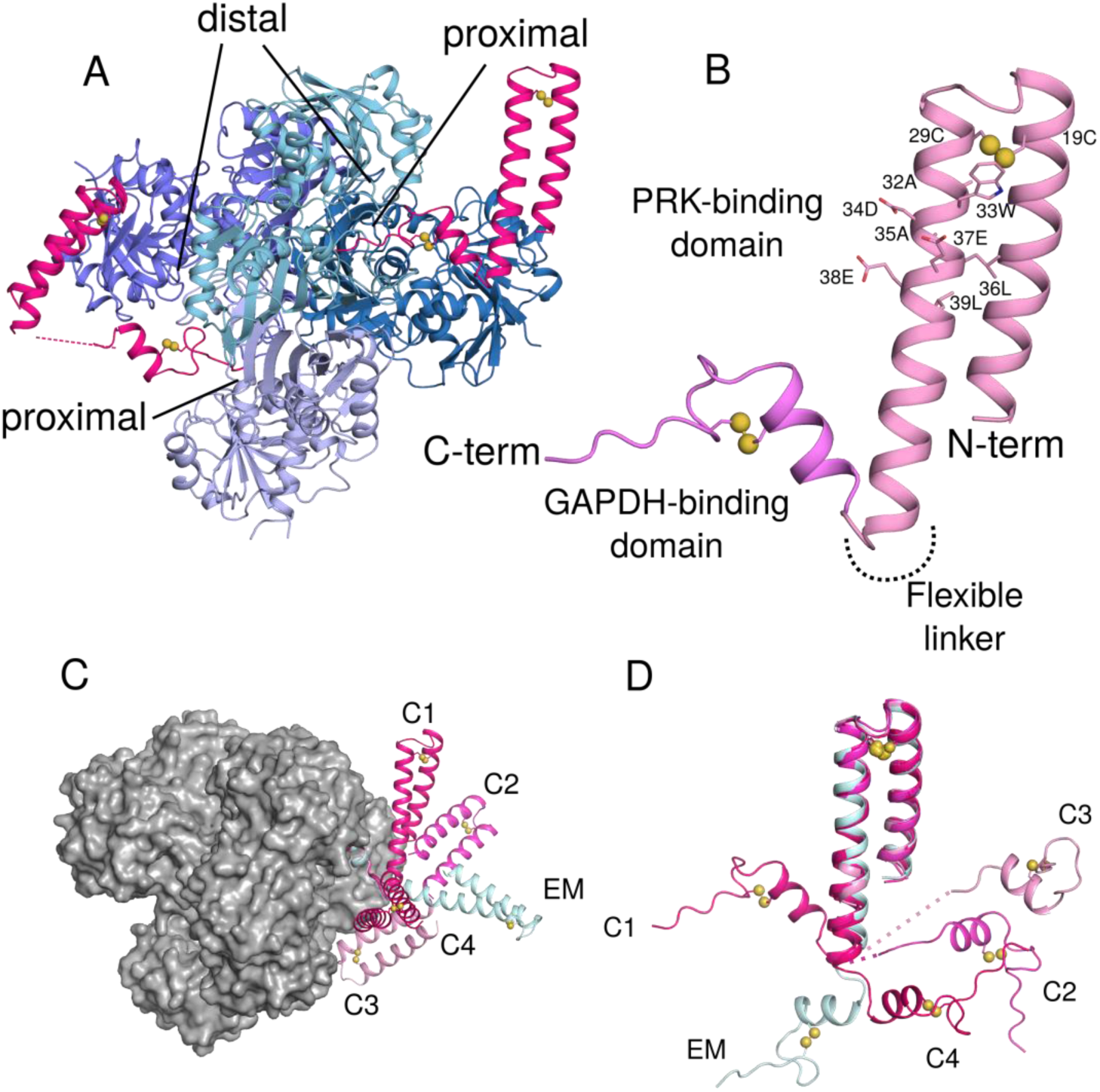
Crystal structures of GAPDH-CP12 complex with full-length CP12. (A) GAPDH tetramer (blue) with two active sites bound by CP12 (pink). Residues 55-75 of CP12 are inserted in the active site of GAPDH, while 1-55 form a two-helix PRK-binding domain (B) CP12 is formed of two domains connected by a flexible linker. The N-terminal PRK binding domain is formed of two anti-parallel helices connected by a disulfide bridge at the apex of the helices. Residues of the conserved CP12 AWDA(V/L)EEL motif are shown as sticks; disulfide bonds are indicated by yellow spheres. The C-terminal GAPDH binding domain is formed of a 2-turn helical region followed by the remaining C-terminal residues that insert into the GAPDH active site. This fold is also stabilized by a disulfide bridge. (C) The different conformations of CP12 observed in crystals (C1-C4) and cryoEM (EM) superposed on the C-terminal region, bound to a GAPDH tetramer (gray surface). (D) CP12 conformations superposed on the N-terminal PRK-binding domain.

To investigate the molecular basis for how CP12 inhibits PRK activity, we solved the structure of the full ternary GAPDH-CP12-PRK complex by cryoEM. We assembled the ternary complex by reconstituting recombinant GAPDH-CP12 with native PRK, partially purified from *T. elongatus.* We enriched PRK activity from cell lysate and incubated it with recombinant GAPDH-CP12 complex to form the ternary complex, which was isolated by size exclusion chromatography (fig. S2-4). Frozen-hydrated samples were imaged in the electron microscope, and used to generate a single-particle reconstruction with D2 symmetry to 3.9 Å (FSC), with local resolution ranging from 3.7 Å to 5.6 Å (Fig. 2A and fig. S5). Our GAPDH-CP12 crystal structure together with a partial model of PRK from an archaeal homolog (*13*) was used as a basis for model building of the entire complex in Coot (*14*), and model refinement with Phenix (*15*) real space refine (Fig.2B, table S2).

**Fig. 2.**
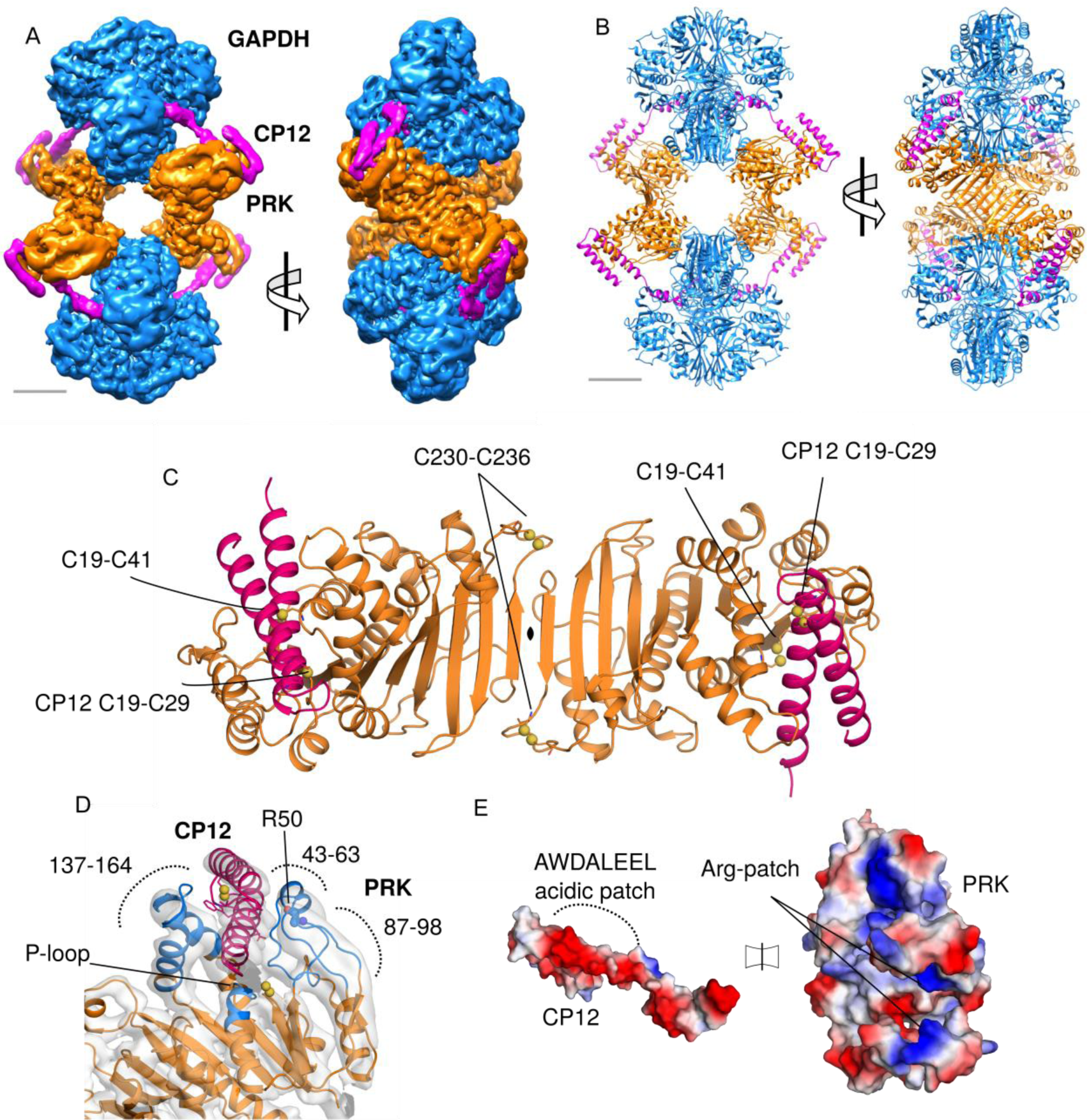
CryoEM reconstruction of GAPDH-CP12-PRK complex. (A) CryoEM maps of the complex with GAPDH (blue), PRK (orange) and CP12 (pink), scale bar 30 Å (B) Cartoon view of the complex with two GAPDH tetramers and two PRK dimers forming an elongated diamond shaped complex tethered by four CP12 chains. (C) Cartoon view of PRK dimer, with CP12 bound in the two active sites; disulfides in both proteins labeled. (D) CP12 (pink) sterically blocks PRK (orange) by binding in the active site (blue). The PRK active site cleft is formed of regions 147-164, 43-63 and 87-98. The PRK ATP-binding P-loop region and Arg50 are indicated. (E) Electrostatic surface potential of CP12 and PRK interface regions, showing charge complementary. The CP12 conserved AWDA(V/L)EEL motif is highlighted.

The inhibited GAPDH-CP12-PRK ternary complex has a hollow diamond-shaped architecture. GAPDH tetramers comprise two vertices separated by 200 Å, while the other two opposite vertices are formed by PRK dimers. Similar to our crystal structures of the binary complex, each GAPDH tetramer binds two copies of CP12. CP12 bridges the active sites of GAPDH and PRK, locking the complex in an inhibited conformation. The four CP12-PRK interaction interfaces were the least well resolved regions of the D2 symmetrized reconstruction (fig. S5). A reconstruction with no symmetry constraints revealed variations in density across the different CP12-PRK interfaces, but all CP12-GAPDH interfaces within the ternary complex remained well ordered and fully occupied. We used 3D classification to enrich for fully occupied CP12-PRK interfaces; however, global flexibility of the assembly may be limiting our ability to improve resolution (fig. S6).

The CP12 N-terminal helical hairpin forms the interface with PRK. It has a main-chain RMSD of 0.8 Å (Fig. 1D) to the conformation present in the GAPDH-CP12 crystal structures and is similar to that observed in the cyanobacterial CP12-CBS domain protein structure (*9*). Our full-length CP12 structures show that the oxidized N-terminal region of cyanobacterial CP12 is ordered prior to binding PRK and oxidation via disulfide bridge formation primes this interaction. Given the similarities of all observed CP12 conformations together with high sequence conservation (fig. S7), we propose that this is regulatory mechanism is evolutionarily conserved across cyanobacteria, algae, and plants.

Green-type PRK is dimeric, with an alpha-beta-alpha sandwich fold where the central nine-strand beta-sheet is continuous across the dimer interface (Fig. 2C). The active site cleft lies between three loops (residues 137-164, 43-63, and 87-98). The N-terminal helical bundle of CP12 plugs this cleft and sterically blocks the active site (Fig. 2D). The charged patch created by the CP12 motif binds to complementary positively charged regions in the PRK active site (Fig. 2E). Variants in the CP12 of *Chlamydomonas reinhardtii,* equivalent to Trp33, Glu27 and Glu38 in the conserved CP12 motif resulted in loss of complex formation (*16*). The conserved Trp33 is on the surface and packs against PRK, contradicting a prediction that it is buried (*17*). In the PRK of *C. reinhardtii*, Arg64 is required for both ternary complex formation, and full activity (*18, 19*). In our structure the equivalent residue, Arg50, is adjacent to the active site and contacts the outer face of the CP12 helical hairpin via the CP12 motif (Fig. 2D). CP12 residues Asp34 and Glu38 from the motif (Fig. 1B) are candidates for forming salt-bridge interactions with Arg50. Green-type PRK has two conserved pairs of cysteine residues that form another tier of redox regulation (*3*). Cys230 and Cys236 form a disulfide bond in the loop at the end of the sheet at the dimer interface (Fig. 2C) and are required for complex formation in *C. reinhardtii* PRK (*3*). Cys19 is in the middle of the ATP-binding Walker A motif (P-loop, residues 15-24, fig S8) and forms a disulfide with Cys41. When this disulfide bond is oxidized in free PRK, up to 20% activity is retained (*9, 20, 21*). The obligate sequential assembly of the inhibited ternary complex is driven by the relative strengths of interaction interfaces within the complex. In the ternary complex, PRK and GAPDH share a relatively small, non-conserved interface (figs. S8, S9) of only 386 Å^2^ buried surface area per GAPDH-PRK pair, with a predicted binding energy of only −3.9 kcal/mol (*22*) (fig. S10). We conclude that the ternary complex formation is dominated by the extensive contacts of GAPDH and PRK with CP12. The GAPDH-PRK interactions may contribute to the stability of the complex, but are not sufficient for complex formation. We tested the stability of the complex using native gel electrophoresis and mass spectrometry after overnight incubation with combinations of NAD^+^, NADPH, ATP, and ADP. NADPH did not dissociate the cyanobacterial complex in our hands, in contrast to reports on other species (*7, 20, 23*). Only reduction of disulfide bonds of CP12 and PRK with DTT (dithiothreitol) reduced the disulfide bonds of CP12 and PRK, and dissociated the ternary complex (fig. S11). Physiologically, the complex is dissociated by reduced thioredoxin, generated by the light reactions (*7*).

CP12 sterically blocks the distal and proximal GAPDH active sites in both the binary and ternary inhibitory complexes. The conserved CP12 Glu69 (fig. S7) (*10, 11*) clashes with the position of the 2’ phosphate group in the distal GAPDH active site (Fig. 3A), rendering NADP(H) and CP12 binding to GAPDH mutually exclusive. To investigate the effect of this clash on nicotinamide dinucleotide specificity and enzymatic activity, we used Michaelis-Menten kinetics to model the activities of GAPDH and PRK in their active and inhibited states (Fig. 3). We assessed PRK activity by measuring ADP production from ATP and ribulose 5-phosphate (Ru5P) in a coupled assay with ADP-hexokinase. As expected, PRK was completely inhibited in the ternary complex, where all the active sites are blocked, and activity was restored after reduction with DTT (Fig. 3B, table S3). GAPDH is a reversible enzyme, catalyzing the reduction of NAD(P)^+^ or oxidation of NAD(P)H. The physiological Calvin cycle reaction is reduction of bisphosphoglycerate (BPG) by NADPH, however the short-lived nature of BPG made it challenging to measure this activity accurately. Therefore, we measured GAPDH kinetics by an *in situ* coupled reaction following NAD^+^ or NADP^+^ reduction by glyceraldehyde 3-phosphate GAP at 340 nm. We found that the apparent K_m_ of GAPDH for NAD+ and NADP+ were similar, as were the apparent k_cat_ values (Fig. 3C, 3D and table S4). There is a small structural shift, mainly of Arg81 (Fig. 3A), in GAPDH depending on whether NAD^+^ or NADP^+^ is bound, however this does not affect activity with either substrate. When CP12 was bound to GAPDH, activity decreased, and the enzyme was more specific for NAD+. The GAPDH-CP12 complex showed a much higher apparent K_m_ for NADP^+^ than NAD^+^, which we attribute to CP12 being a competitive inhibitor for NADP^+^. The GAPDH-CP12 complex retained a high affinity for NAD^+^, although the rate of turnover was slower. These data could be explained by CP12 blocking the proximal GAPDH active sites and the steric hindrance of CP12 slowing NAD(H) binding to the remaining distal active sites. In the ternary complex, GAPDH activity with NAD^+^ was similar to the binary complex, implying that NAD^+^ can still access the open CP12 distal sites as in the GAPDH-CP12 binary complex. In contrast, ternary complex activity with NADP^+^ was not measurable. These data suggest that cooperative avidity effects of ternary complex formation prevent NADP^+^ from dissociating CP12, which is bound more tightly in the ternary complex. This agrees with the dissociation experiments, where the complex was disassembled only by reduction with DTT and not by NADPH (fig. S11). The differential behavior of the complex to NADPH and NADH, where activity with the physiological substrate NADPH is more inhibited, shows that the complex function integrates responses from both redox state and nicotinamide dinucleotide availability.

**Fig. 3.**
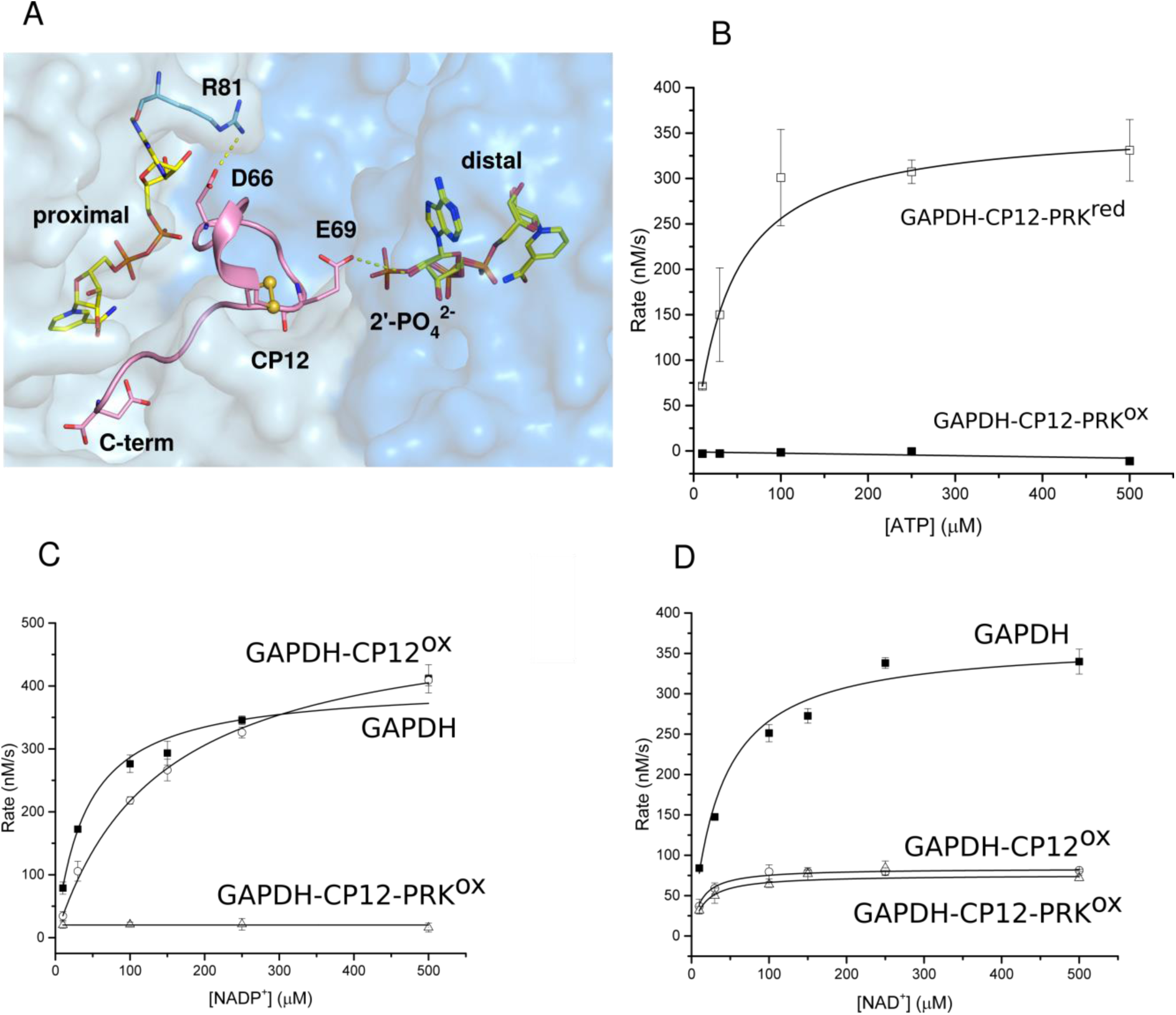
Kinetics and co-enzyme binding of GAPDH complexes. (A) View of GAPDH (NAD^+^)-CP12 with adjacent active sites. The CP12-bound (proximal) site makes extensive contacts with both NAD^+^ and GAPDH. CP12-Glu69 prevents clashes with NADPH binding to the distal site, as it would clash with the 2’-phosphate. (B) PRK activity for oxidized (■) and reduced (□) GAPDH-CP12-PRK complex. All reactions were measured in triplicate and fitted using Michaelis-Menten kinetics. Rate of NADP^+^ reduction (C) and NAD^+^ reduction (D) were measured for GAPDH (■), GAPDH-CP12 (○) and GAPDH-CP12-PRK (r) complexes at increasing concentrations of nucleotide substrate. The maximal velocity (V_max_) with NADP^+^ is not inhibited in GAPDH-CP12 but is fully inhibited in GAPDH-CP12-PRK. In contrast, the V_max_ with NAD^+^ are equally inhibited by GAPDH-CP12 and GAPDH-CP12-PRK.

We investigated how structural changes in CP12 regulate substrate availability for carbon fixation in the Calvin cycle. We combined X-ray crystallography and single-particle cryoEM to determine the structures of two regulatory complexes. Our data provide a mechanism for how GAPDH and PRK activities are redox-regulated in response to light (Fig. 4). CP12 is a conditionally disordered protein (*24*), where a redox-induced change causes a functional and structural switch. When bound to GAPDH, CP12 forms a disulfide-locked helical hairpin which is ordered before interaction with PRK. GAPDH-CP12 captures PRK dimers to form a ternary complex that restricts production of the substrate for Rubisco and prevents reduction of carboxylic acids to sugar. Disulfide bonds within PRK and CP12 are oxidized in the dark and maintain the inhibited complex. PRK binding further increases the stability of the GAPDH-CP12 complex, thus restricting NADPH turnover. Our observations provide a redox-sensitive molecular mechanism that controls the “off switch” for how plants make biomass.

**Fig. 4.**
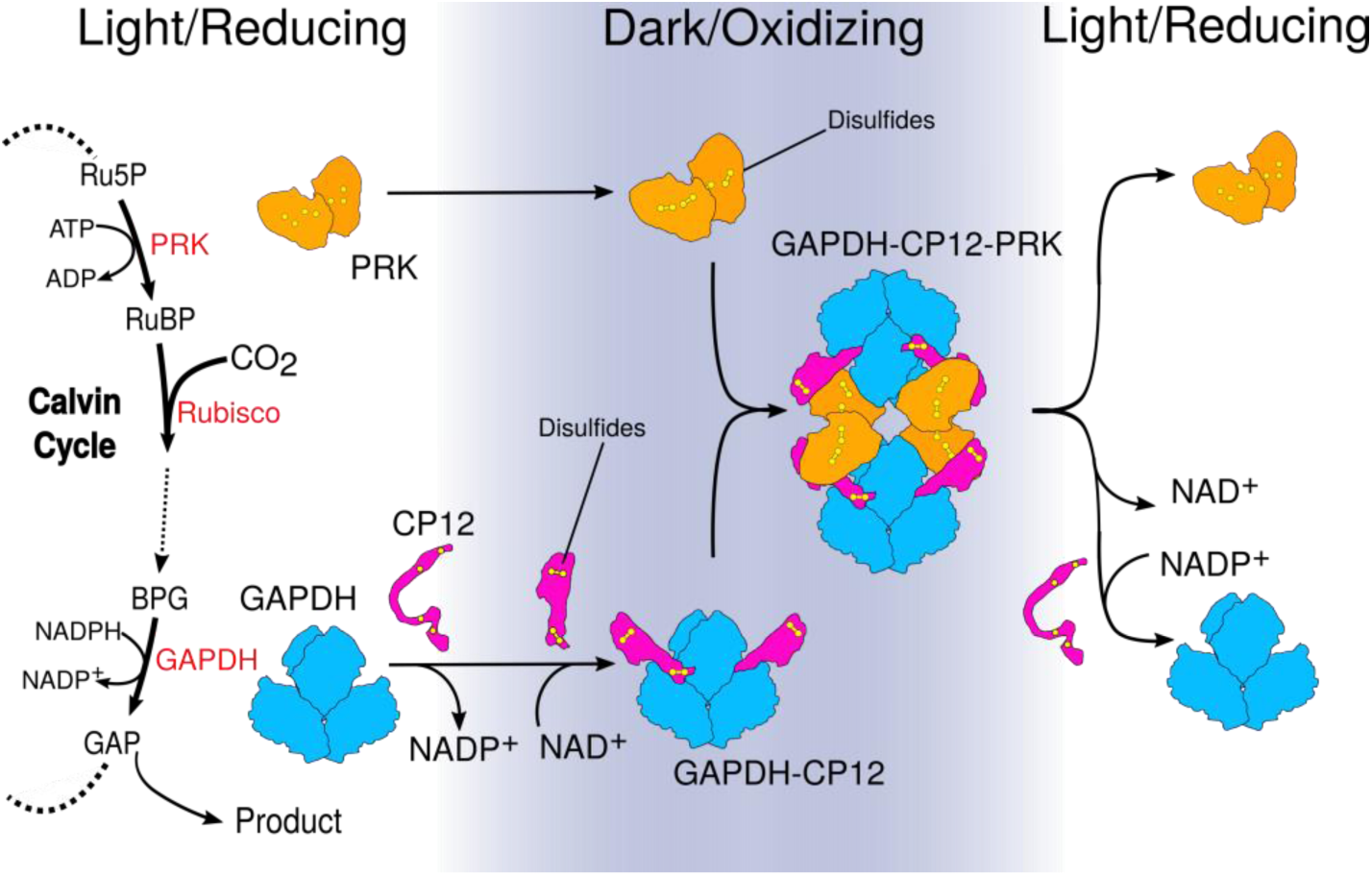
Model for how structural changes in redox-dependent complexes regulate carbon fixation. GAPDH and PRK are energy consuming enzymes of the Calvin cycle. The PRK catalyzed step directly precedes Rubisco carbon fixation. GAPDH (blue) activity is at the branch point between regeneration of RuBP or central metabolism. In the dark, the oxidizing environment causes intramolecular disulfide bridges to form within CP12 (pink) and PRK (orange). In parallel NADP^+^ bound to GAPDH is exchanged for NAD^+^. CP12 binds to GAPDH, reducing its activity. Pre-ordered CP12s subsequently recruit PRK, blocking PRK active sites to substrate and GAPDH active sites to NADP(H). When returning to the light, disulfide reduction by thioredoxin dissociates the complex, releasing GAPDH and PRK.

## MATERIALS AND METHODS

### GAPDH and CP12 expression and purification

The GAPDH (*tll1466*) and CP12 genes (*tsr1820*) were amplified by PCR from *Thermosynechococcus elongatus* BP-1 (a gift from Matthias Rögner, Rühr-Universität Bochum) genomic DNA (primers CP12-FOR

TTCCGCGTGGATCCATGAGTAATCTCGAG; CP12-REV GCTGCAGATCTCGAGTTATTAGTCGTCG; GAPDH-FOR GTCCGCGTGGATCCATGGTTAGAGTCGC; GAPDH-REV

GCCGGATCCTCGAGCTACTAAGCCCAGTGG), and cloned by restriction-ligation into a modified prSET-A expression vector (*25*) with a thrombin-cleavable hexahistidine tag.

*Escherichia coli* KRX (Promega) transformed with the GAPDH or CP12 plasmids were grown in Terrific Broth with 100 μg/ml ampicillin at 37°C to OD _600_ 0.6-0.8 followed by overnight induction at 18°C with 0.1 % (w/v) rhamnose. Cells were harvested by centrifugation, resuspended in 50 mM Tris-HCl, 150 mM NaCl, pH 7.9 and lysed by sonication. Insoluble material was removed by centrifugation for 1 hour at 80,000 g. Protein was purified by Ni-NTA affinity chromatography. The 50 ml of lysate supernatant was incubated with 3 ml of Ni-NTA resin (Qiagen). The resin was washed three times with 15 ml of 50 mM Tris-HCl, 150 mM NaCl, 30 mM imidazole followed by elution in the same buffer with 300 mM imidazole.

The eluted proteins were cleaved overnight at 4°C with 100 units of thrombin (Sigma-Aldrich), concentrated to 1 ml (Vivaspin concentrators, Sartorius) and further purified by size-exclusion chromatography (Proteosec 6-600 column, Generon). The flow rate was 1 ml/min (Äkta Purifier) with a mobile phase of 50 mM Tris-HCl, 150 mM NaCl, pH 7.9. The GAPDH-CP12 complex was made by mixing with a molar ration of 10 CP12 per GAPDH monomer, followed by size-exclusion chromatography.

### Purification of ternary complex

PRK was purified from *T. elongatus* cell lysate. Throughout the purification, PRK-containing fractions were identified by assaying for PRK activity. *T. elongatus* was grown in batches of 3 l in DTN media (*26*) and 20 l were used in a preparation. Cultures were grown to OD_720_ 1.0 at 45°C under constant illumination at 20 μmol m^-2^ s^-1^. Cells were pelleted and washed in 50 mM Tris-HCl, 150 mM NaCl, 10 mM MgCl_2_, pH 7.9 and lysed with a cell disruptor at 25 kpsi (Constant systems T5) followed by centrifugation for 1 hour at 185,000 g.

The lysate supernatant was dialyzed for 16 hours against 50 mM Tris-HCl pH 7.9. The dialyzed sample was 0.22 μm filtered and loaded on DEAE-s (Toyopearl) 75 ml anion exchange column using a peristaltic pump at 2 ml/min (P-1 GE Healthcare). Protein was eluted with an NaCl gradient from 0 to 150 mM over seven column volumes (Äkta Purifier). The anion exchange fractions with PRK activity were dialyzed for 16 hours against 10 mM phosphate buffer pH 6.5. Protein was then loaded onto a 35 ml hydroxyapatite column (Bio-rad) using a peristaltic pump at 2 ml/min (P-1 GE Healthcare). Protein was eluted using a gradient to 250 mM phosphate buffer pH 7.0 over five column volumes.

PRK-containing fractions from the hydroxyapatite elution were then further purified using size-exclusion chromatography (Proteosec 6-600, Generon). This was carried out with a flow rate of 0.5 ml/min and a mobile phase of 50 mM Tris-HCl, 150 mM NaCl, 10 mM MgCl_2_, pH 7.9. The partially purified PRK was incubated overnight with a 5 molar excess of GAPDH-CP12 binary complex with 1 mM NAD^+^, 1 mM trans-1,2-dihydroxy-4,5-dithane (oxidized DTT) and 1 mM ADP (Sigma-Aldrich). The ternary complex was purified using size-exclusion chromatography (Proteosec 6-600 column, Generon). This was carried out with a flow rate of 0.5 ml/min (ÄKTA Purifier) and a mobile phase of 50 mM Tris-HCl,50 mM NaCl, 10 mM MgCl_2_, pH 7.9. The eluted complex was identified using SDS-PAGE and ESI-QUAD-TOF mass spectroscopy (*27*).

Protein concentration was determined using Pierce-BCA assays (ThermoFischer).

### Enzyme activity assays

GAPDH activity was measured by following the reduction of NAD(P)^+^ at 340 nm with DL-glyceraldehyde-3-phosphate and arsenate. Each 200 μl reaction contained 5 Mm DL-glyceraldehyde-3-phosphate (Sigma-Aldrich), 2.5 mM sodium arsenate (Sigma-Aldrich) in 10 mM Tris, 50 mM NaCl, 10 mM MgCl_2_, pH 7.9. GAPDH activity to measure the oxidation of NAD(P)H was measured using a coupled enzyme that produced the GAPDH substrate 1,3-bisphosphoglycerate. Each 200 μl reaction contained 5 mM 3-phosphoglycerate, 5 mM ATP and 1 unit of yeast phosphoglycerate kinase (Sigma) in 10 mM Tris, 50 mM NaCl, 10 mM MgCl_2_, pH 7.9. Reactions were performed at 25 °C in a quartz cuvette (Hellma) and absorbance at 340 nm (A340) was measured using a 1 nm slit-length and 1 Hz sampling rate (Shimadzu).

PRK activity was measured by coupling the production of ADP product to the reduction of NAD^+^ using ADP-hexokinase and glucose-6-phosphate dehydrogenase (fig. S2A) and following increase in A340. In each reaction, the coupling enzyme reagent contained 10 μg/ml ADP-hexokinase, 4 units of glucose-6-phosphate dehydrogenase (Sigma-Aldrich), 10 mM glucose and 500 μM NAD+ in 10 mM Tris, 50 mM NaCl, 10 mM MgCl_2_, pH 7.9. Reducing conditions for assays were achieved by pre-incubation of the sample for 15 min with 10 mM DTT.

The ADP-dependent hexokinase gene (WP_004069859) from *Thermococcus litoralis* DSM-5473 gDNA (Deutsche Sammlung von Mikroorganismen und Zellkulturen) (*28*) was PCR amplified and cloned by Gibson assembly into the modified pRSETA vector and purified by affinity chromatography as for GAPDH and CP12.

For the purification of PRK, activity was measured at 25 °C in a 150 μl reaction containing the coupled enzyme assay mix with 5 μl of protein sample. Activity was measured with 100 μM ATP plus 100 μM ribulose-5-phosphate (Ru5P, Sigma-Aldrich) corrected for background ATPase activity by measuring the same sample with 100 μM ATP and no Ru5P (fig. S3B).

Michaelis-Menten kinetics for both GAPDH and PRK were determined by measuring the initial reaction rates for 30 s at increasing concentrations of substrate in the presence of the equivalent 55 nmols of GAPDH monomer and 25 nmols PRK monomer. Traces were plotted and analyzed using Origin (OriginLab). Each trace was measured for 100 s with the initial rate measured by fitting the first 30 s to a linear curve using a least-squares function. Activity at each substrate concentration was measured in triplicate and plotted with standard deviation.

### Native-gel electrophoresis

Native-gel PAGE was performed on GAPDH-CP12-PRK samples by incubating 20 μl of 100 μg/ml aliquots overnight in combinations of DTT, NADPH, ADP, each at 2.5 mM. Samples were loaded onto 4-20 % gradient Tris-Glycine gels (NuPAGE, Invitrogen). The loading buffer contained 62.5 mM Tris-HCl, 0.05 % bromophenol blue, 10 % glycerol, pH 6.8. The running buffer was 25 mM Tris base, 192 mM glycine. Gels were loaded and run on an XCell SureLock Mini-Cell electrophoresis system (Thermo-Fischer) at constant-current set at 20 mA for 4 hours followed by staining with Instant Blue Coomassie (Expedion). Protein bands were identified using ESI-QUAD-TOF mass spectroscopy performed by the St. Andrews mass spectrometry service (*27*)

### Protein crystallization

GAPDH and GAPDH-CP12 complexes at 50 mg/ml were screened for crystallization with sitting drop vapor diffusion at 17°C using commercial sparse matrix screens JCSG-plus, Wizard 3 and 4 (Molecular Dimensions), PEG ION 1 and 2 (Hampton) commercial screens with a Mosquito robot (TTPLabTech). Some hits were further optimized in manually set up hanging drop vapor diffusion experiments. For the structure of GAPDH with four CP12 bound, GAPDH-CP12 was incubated with a further 10-fold molar excess of CP12 to GAPDH monomer. The structure of GAPDH with NADPH bound was obtained by mixing GAPDH with 5 mM of glyceraldehyde-3-phosphate, 3-phosphoglycerate, NADP^+^, NADPH and 1 unit/ml bisphosphoglycerate kinase prior to crystallization turning over the bound NAD^+^ in GAPDH, replacing it with NADP^+^. Crystallization conditions are given in Table S1. Crystals were cryo-protected in the mother liquor with 30% volume PEG 400 added, then flash-cooled in liquid nitrogen.

### X-ray structure determination

X-ray diffraction data were collected at Diamond Light Source synchrotron, and processed with the xia2 pipeline (*29*). GAPDH crystal structures were solved by molecular replacement in Phaser (*30*) with a model based on our earlier *T. elongatus* GAPDH structure (PDB 4BOY). The models were rebuilt in Coot (*14*) with cycles of refinement in phenix.refine (*31*), and validated with MolProbity (*32*). The CP12 models could be built into the difference density. Data collection and refinement information are in Table S1. Coordinates and structure factors were deposited in the PDB with accessions 6GFP, 6GFQ, 6GFR, 6GFO, 6GG7, 6GHR, 6GHL.

Structure figures were prepared using PyMol (Shrödinger) and UCSF Chimera (*33*).

### Negative stain EM

Negative stain EM was used to assess sample quality during purification. 2.5 μl of the GAPDH-CP12-PRK complex was applied to glow-discharged carbon-coated copper grids (400 mesh) and stained with 2% uranyl acetate. Data were collected on a 120 keV Tecnai T12 microscope with a pixel size of 2.6 Å.

### Sample preparation and cryoEM data acquisition

Immediately following size-exclusion chromatography, 2.5 μl GAPDH-CP12-PRK complex in 50 mM Tris-HCl, 30 mM NaCl, 10 mM MgCl_2_, 1 mM NAD^+^, 1 mM ADP, 1 mM 1 mM trans-1,2-dihydroxy-4,5-dithane was adsorbed to a glow-discharged holey carbon grid (C-Flat 1.2/1.3 grid or Quantifoil R2/2), which was overlayed with a thin layer of amorphous carbon. Using a Vitrobot mark III (Thermo Fisher Scientific), grids were blotted for 3.5 seconds at “blot force” 3 and plunge frozen in liquid ethane cooled to liquid nitrogen temperature. For initial model generation, data were collected on a 200 keV Tecnai F20 electron microscope (Thermo Fisher Scientific) fitted with a Falcon II direct electron detector (Thermo Fisher Scientific). Images were recorded at a pixel size of 2.05 Å and with a defocus range of 4 to 6 μm underfocus. Electron micrograph movies for the high resolution reconstruction were collected on a 300 keV Titan Krios (Thermo Fisher Scientific) equipped with a Quantum K2 Summit direct electron detector (Gatan). Data were collected in counting mode, and image stacks were recorded at 6 frames per second for 9 seconds with an accumulated dose of 44 e^-^/Å^2^. Images were recorded at pixel size of 1.047 Å and with a defocus range of −1.75 to −3.5 μm. All data were acquired using image acquisition software EPU (Thermo Fisher Scientific).

### CryoEM image processing

An initial model was generated using data collected at 200 keV on a Tecnai F20 electron microscope. Contrast transfer function (CTF) parameters were estimated using CTFFIND4 (*34*), and micrographs were curated based on figure of merit value and ice quality. 14,000 particles were manually picked from the remaining micrographs, and 2,708 particles were discarded after 2D classification and selection. The initial model generation tool within RELION (*35*) was used to create a model which was refined with C2 symmetry constraints to a resolution of 17.4 Å. Additional rounds of 2D classification enabled the selection of a more homogenous subset, and the remaining 5,828 particles were used to refine the particle orientations with D2 symmetry constraints. The resulting 11.2 Å resolution reconstruction was then used as an initial model for the refinement of Krios-collected data.

For the high resolution cryoEM reconstruction, 4880 micrograph movies were collected at 300 keV on the Titan Krios electron microscope. Electron micrograph movie frames were aligned by MotionCor2 (*36*), discarding the first and last frames. CTF parameters were estimated using CTFFIND4 (*34*). Any movies containing low figure of merit scores, substantial drift, low contrast, thick ice, or crystalline ice were discarded from further analysis. 2D class averages from cryoEM data were used as templates for the automated particle picking tool in RELION. 585,526 initial particles were subjected to multiple rounds of 2D classification and selection. The initial model derived from data collected at 200 keV was strongly low-pass filtered (40 Å) to prevent model bias and used as a starting model for a gold-standard 3D autorefinement of images. These orientations served as the starting point for tracking beam-induced movement of individual particles, which was corrected using particle polishing within RELION. Additional rounds of 2D classification after particle polishing were performed to improve the sample homogeneity. 197,212 selected particles contributed to a final reconstruction which was refined using D2 symmetry. The overall resolution of 3.9 Å was calculated using the mask-corrected Fourier shell correlation (FSC) with local resolution ranging from 3.7 to 5.6 Å (fig S5). The final reconstruction was deposited in the EMDB with accession code EMD-0071. Handedness of the reconstruction was determined by fitting the chiral GAPDH crystal structure into the 3.9 Å map using the ‘fit in map’ tool in Chimera (*33*). There was a clear difference between correlation coefficients when comparing the different hands (CC = 0.86 versus 0.65).

To assess the extent of conformational and stoichiometric heterogeneity of the GAPDH-CP12-PRK complex, we also performed a refinement without symmetry constraints. The resulting reconstruction refined to 4.3 Å and revealed heterogeneity across the four CP12-PRK binding sites. We implemented a number of 3D classification strategies within the RELION framework in an attempt to select for homogenous complexes, however none improved the overall resolution or local resolutions of the reconstructions. In one strategy, particles were classified into 6 groups allowing orientations to be refined after each iteration. The two most populated classes, which also reported the highest resolution estimates, were combined and used in a 3D autorefinement imposing either C1 or D2 symmetry. In the second strategy, the densities for GAPDH and the regions of PRK that do not interact with CP12 were subtracted from each particle. Density-subtracted particles were separated into 3 classes using 3D classification without refinement, focusing on the least occupied CP12-PRK interface.

### CryoEM model bBuilding and refinement

The crystallographic GAPDH-CP12 model was placed in the B-factor sharpened, local resolution-filtered map with Chimera, and the CP12 N-terminus and linker was refitted into the density. A PRK model based on an archaeal PRK (PDB 5B3F)(*13*) was placed in the map and extensively rebuilt and extended with the *T. elongatus* sequence. The asymmetric unit was rebuilt in COOT (*14*), expanded to complete the D2 symmetry and refined with phenix real space refine (*15*) using rotamer restraints and secondary structure restraints derived from the crystallographic models. The final model was validated with MolProbity (*32*). A locally sharpened map made with phenix.auto_sharpen (*37*) was also used in model building. The final model is full-length in the three proteins, however side chains for less well ordered regions were omitted in the final coordinates, with most missing from the N-terminal regions of PRK and CP12. Parameters for the cryoEM ternary complex model are given in table S2. Coordinates were deposited in the PDB with accessions, 6GVE.

## ACKNOWLEDGEMENTS

We thank Diamond for access and support of the CryoEM facilities at the UK national electron bio-imaging center (eBIC), (proposals EM19432 and EM18659), funded by the Wellcome Trust, MRC and BBSRC. *We thank Diamond Light Source for X-ray beam time (proposal mx12579), and the staff of beam lines I02, I03, I24, and I04 for assistance with crystal testing and data collection.* This work was supported by a BBSRC Doctoral Training Programme grant (BB/J014575/1 to C.M.). We thank project students Nishat Miah and Laura Briggs. C.M., and N.S., performed the experiments. Data were analyzed by C.M., N.S., D.B. and J.W.M. B.V.K. and C.A.R.C. did cloning and early biochemical and structural work. The paper was written by J.W.M., C.M., N.S. and D.B.. All authors approve of the conclusions and final version of the paper. The atomic coordinates and structure factors for GAPDH-NAD, GAPDH-NADP, GAPDH-CP12-conf1-conf2, GAPDH-CP12-conf3, GAPDH-CP12-conf4, GAPDH-CP122 and GAPDH-CP12_4_ have been deposited in the Protein Data Bank under accession codes 6GFR, 6GFP, 6GFO, 6GHR, 6GHL, 6GFQ and 6GG7 respectively. The GAPDH-CP12-PRK cryoEM map has been deposited in the Electron Microscopy Databank under EMDB-0071, and the atomic model has been deposited in the PDB, accession 6GVE. Structural statistics are provided in the supplementary materials (tables S2 and S3). The authors declare no competing financial interests.

